# BMSC-Laden PEGDA/HAMA Dual-Crosslinked Hydrogels with Distinct Stiffness Profiles for Osteochondral Defect Repair

**DOI:** 10.64898/2026.05.24.727447

**Authors:** MaoJiang Lyu, Xin Guo, LiQi Ng, Yi Sun, Jianjing Lin, XiaoFeng Zhang, XinTao Zhang, YaoHua He

**Author notes:** Corresponding author.: Email addresses (X. Zhang). Corresponding author. Department of Orthopedic Surgery, Shanghai Sixth People’s Hospital Affiliated to Shanghai Jiao Tong University School of Medicine, Shanghai, China. Email addresses (Y. He). These authors contributed equally (MaoJiang Lyu, Xin Guo, LiQi Ng).

## Abstract

This study aimed to develop a BMSC-laden polyethylene glycol diacrylate/methacrylated hyaluronic acid (PEGDA/HAMA) dual-crosslinked hydrogel and evaluate its effects on osteochondral defect repair. Two PEGDA concentrations, 3.75% and 7.5% (w/v), were used to prepare representative soft and stiff hydrogel formulations, respectively. The hydrogels were characterized in terms of morphology, cytocompatibility, and compressive behavior, and their ability to support BMSC-associated matrix deposition was evaluated in vitro. Repair outcomes were further assessed in a rat osteochondral defect model at 4 and 8 weeks. The 7.5% PEGDA/HAMA hydrogel exhibited higher stiffness than the 3.75% formulation and supported BMSC viability and matrix deposition in vitro. In vivo, the stiff hydrogel group showed improved defect filling and subchondral bone remodeling compared with the soft hydrogel and defect groups. However, histological and immunohistochemical analyses revealed predominant collagen type I deposition and limited collagen type II expression in the repair region, indicating fibrocartilaginous rather than hyaline-like cartilage repair. These findings suggest that BMSC-laden PEGDA/HAMA hydrogels may provide a useful platform for osteochondral defect repair, while further optimization of degradation behavior, matrix maturation, and collagen type II deposition is required to improve hyaline cartilage-oriented repair.

## Introduction

Articular cartilage injuries represent a prevalent and challenging clinical problem, with epidemiological studies revealing cartilage lesions in over 60% of knee arthroscopies [1,2]. Due to the inherently limited self-repair capacity of cartilage tissue, such injuries often progress to osteoarthritis without effective intervention [3-5]. Current standard clinical treatments, including microfracture, can stimulate mesenchymal stem cell migration to the defect site but primarily generate mechanically inferior fibrocartilage rather than native hyaline cartilage, resulting in suboptimal long-term outcomes [6,7].

Tissue engineering offers a promising alternative strategy for cartilage regeneration, primarily utilizing biodegradable scaffolds as cell carriers and provisional matrices [8-11]. Among these, hydrogels are considered ideal scaffold materials due to their high water content, which resembles the native extracellular matrix, their excellent biocompatibility, and their capacity for minimally invasive injection with in situ formation [12,13]. An ideal cartilage repair scaffold must fulfill two critical requirements: sufficient early-stage mechanical strength to withstand dynamic joint loading and maintain defect filling stability, and seamless integration with host tissue, including osseointegration with subchondral bone and interfacial integration with surrounding cartilage [14-18].

Hyaluronic acid (HA), a natural component of the extracellular matrix, has emerged as a promising biomaterial for cartilage tissue engineering due to its inherent bioactivity, cell-mediated enzymatic degradability, and excellent chemical modifiability [19,20]. However, pure HA hydrogels exhibit weak mechanical strength and rapid degradation, limiting their standalone application [21,22]. In contrast, synthetic polyethylene glycol (PEG)-based hydrogels offer precise control over mechanical properties and degradation kinetics but lack essential bio-recognition signals [23,24]. Therefore, developing HA/PEG hybrid hydrogels that combine the respective advantages of both has become an important research direction [19,23-24].

Although PEG- and HA-based hydrogels have been widely investigated for cartilage tissue engineering, balancing mechanical stability, cell compatibility, and matrix deposition remains challenging [11,19,23–26]. In particular, the influence of PEGDA content on the mechanical behavior and biological performance of BMSC-laden PEGDA/HAMA hydrogels requires further clarification. Given that insufficient scaffold stiffness may compromise early defect filling and structural stability, whereas excessive stiffness may restrict cell-mediated remodeling, comparing hydrogel formulations with distinct stiffness profiles may provide useful information for optimizing osteochondral repair scaffolds [14–18].

In this study, we developed a BMSC-laden PEGDA/HAMA dual-crosslinked hydrogel system and evaluated its stiffness-modulated effects on osteochondral defect repair. Two PEGDA concentrations, 3.75% and 7.5% (w/v), were selected to establish representative soft and stiff hydrogel formulations, respectively. This binary stiffness-modulated comparison was designed to assess whether increasing PEGDA content could enhance hydrogel stiffness while maintaining injectable handling, photocrosslinking feasibility, and BMSC encapsulation compatibility. The study therefore focused on determining how these two mechanically distinct PEGDA/HAMA hydrogel formulations influenced BMSC-associated matrix deposition in vitro and osteochondral defect repair in vivo.

## Materials and methods

### 2.1 Synthesis of Methacrylated Hyaluronic Acid (HAMA)

HA and PEGDA were purchased from Guangzhou Tanshui Biotechnology Co., Ltd. (Guangzhou, China). HA powder (1 g, 50 kDa) was dissolved in 50 mL of deionized water under continuous stirring at 4 °C overnight. Subsequently, 33.33 mL of dimethylformamide (DMF) was gradually added to the HA solution. MMA (7.5 mmol) was then added dropwise to the mixture under continuous stirring. During MMA addition, the pH was maintained at 8–9 using 1 M NaOH. The reaction was continued at 4 °C for 24 h and then terminated by adding 2.47 g of NaCl. The product was precipitated with ethanol three times, washed, redissolved in deionized water, and dialyzed against deionized water for 3 days using dialysis tubing with a molecular weight cutoff of 3.5 kDa. The final HAMA product was obtained by freeze-drying. The chemical structure of HAMA was analyzed by ¹H NMR.

### 2.2 Preparation of HA/PEGDA Composite Hydrogels

HAMA powder (30 mg, 50 kDa) was dissolved in 1 mL of phosphate-buffered saline (PBS). PEGDA solutions were prepared by dissolving 37.5 mg or 75 mg of PEGDA (10 kDa) in 1 mL of PBS. Lithium phenyl-2,4,6-trimethylbenzoylphosphinate (LAP) was added to achieve a final concentration of 0.25% (w/v) for photoinitiation. The prepared solutions were mixed to obtain final PEGDA concentrations of 3.75% and 7.5% (w/v). The hydrogel precursor solutions were exposed to 405 nm blue light at 50 mW/cm² for photopolymerization. Hydrogel formation was assessed using the inverted vial method [27], and the gelation time was recorded as the time point at which the precursor solution ceased to flow upon vial inversion. The 3.75% and 7.5% PEGDA formulations were selected as representative soft and stiff hydrogel conditions, respectively, to enable a binary stiffness comparison of PEGDA/HAMA hydrogels.

### 2.3 Cell viabilities of photo-cured PEGDA/HAMA hydrogels

SD rat bone marrow mesenchymal stem cells (BMSCs) were obtained from Cyagen (Guangzhou) Biotechnology Co., Ltd. (Guangzhou, China; product number: RASMX-01001). BMSCs were encapsulated within two different hydrogel concentrations during photo-polymerization. For the cell viability test, BMSCs (2.0×10^6^ cells/mL) were combined with each PEGDA/HAMA hydrogel formulation in culture media, followed by photo-crosslinking under 405 nm blue light for 2 minutes. The resulting cell-laden hydrogels were transferred into separate wells of a 24-well tissue culture plate. To each well, 500 μL of fresh DMEM containing 10% FBS was added. The plates were then incubated at 37 °C in a humidified atmosphere with 5% CO_2_ for either 7 or 21 days. Cell viability and proliferation were assessed using a live/dead assay (Life Technologies, NY, USA), and the stained cells were examined using confocal laser scanning microscopy (CLSM, Eclipse E600W, Tokyo, Japan).

### 2.4 Mechanical Evaluation of Hydrogels

An in-situ mechanical testing machine (Ke’er Control & Testing System (Tianjin) Co., Ltd.) was employed to measure the compressive stress–strain curves of the hydrogels. Hydrogels were prepared as cylinders with a diameter of 7 mm and a height of 6 mm. The crosshead at the top was then moved upward at a rate of 50 mm/min until the hydrogel fractured or the machine reached its operational limit. Each test was conducted three times.

### 2.5 Chondrogenic Matrix Deposition in 3D PEGDA/HAMA Hydrogels

To evaluate the chondrogenic potential of PEGDA/HAMA hydrogels, BMSCs at a density of 2.0 × 10L cells/mL were encapsulated within PEGDA/HAMA hydrogel precursor solutions and photocrosslinked using 405 nm blue light in 24-well Transwell inserts (5 mm diameter × 5 mm height, n=6 per group). The constructs were initially maintained in growth medium for 7 days to ensure cell viability and matrix adaptation, followed by chondrogenic induction using differentiation medium for an additional 7 days (total culture period = 14 days). Matrix deposition was evaluated by Safranin O and Masson’s trichrome staining. This assay was designed to evaluate early chondrogenic matrix accumulation rather than mature hyaline cartilage formation.

### 2.6 Scanning Electron Microscope (SEM) Morphology Observation

After lyophilization, the scaffold specimens’ cross-sections were coated with gold and examined using SEM to analyze their morphology and pore sizes.

### 2.7 Rat Osteochondral Defect Model and Hydrogel Implantation

All animal procedures were approved by the Institutional Animal Care and Use Committee of Peking University Shenzhen Hospital under protocol No. 2023-139, approved on 21 June 2023. Male Sprague-Dawley rats weighing 280–300 g were obtained from Guangzhou Jinwei Biotechnology Co., Ltd. and housed in the animal facility of Peking University Shenzhen Hospital. Animals were maintained under a 12 h light/dark cycle at 24 °C and 50–55% relative humidity, with free access to food and water. Rats were randomly assigned to three experimental groups (n = 5 per group): (1) Defect group, (2) Soft hydrogel group, and (3) Stiff hydrogel group. Rats were anesthetized with isoflurane, and the surgical area was shaved and disinfected before surgery. Animals were monitored daily for wound healing, mobility, and signs of distress. A midline incision was made anterior to the knee joint, and an osteochondral defect measuring 2 mm in diameter and 1 mm in depth was created centrally in the femoral trochlea. Immediately after defect formation, 50 µL of the designated treatment was injected into the defect site. The Defect group received phosphate-buffered saline (PBS). For the hydrogel-treated groups, BMSCs were suspended in the PEGDA/HAMA precursor solutions before implantation. The Soft hydrogel group received 50 µL of BMSC-laden 3.75% PEGDA/HAMA precursor solution, whereas the Stiff hydrogel group received 50 µL of BMSC-laden 7.5% PEGDA/HAMA precursor solution. The final BMSC density was 2.0 × 10L cells/mL. After injection, the hydrogel precursor solutions were photocrosslinked in situ using 405 nm light at 50 mW/cm² for 30 s.

At 4 and 8 weeks post-surgery, the rats were euthanized, and the distal femurs, including the defect area and adjacent tissues, were collected. The tissues were fixed in 4% (w/v) paraformaldehyde at 4°C for 72 hours, then decalcified in 10% EDTA at 4°C for one month. Following decalcification, the samples were embedded in paraffin and subjected to routine histological processing.

### 2.8 Gross morphology observation and ICRS scoring

At 4 and 8 weeks after surgery, the distal femurs were harvested and photographed to assess gross defect repair. Gross repair outcomes were evaluated using the International Cartilage Repair Society (ICRS) macroscopic scoring system [28]. Scoring was performed based on defect filling, integration with adjacent tissue, and surface appearance.

### 2.9 Micro-CT analysis

Femur samples with defects were imaged using a micro-CT scanner at a voxel size of 7 μm. From the central defect region, 200 slices were chosen for detailed evaluation. Three-dimensional reconstructions were performed with DATAVIEWER (version 1.5.6.2) and CTvox (version 3.3), while CTAn (version 1.18.4.0) was employed to quantify parameters such as bone volume fraction (BV/TV), trabecular number (Tb.N), trabecular separation (Tb.Sp), and trabecular thickness (Tb.Th).

### 2.10 Histological and immunohistochemical staining assessment

After fixation and decalcification as described above, the distal femur specimens containing the osteochondral defect region were dehydrated through a graded ethanol series, cleared in xylene, and embedded in paraffin. Paraffin sections with a thickness of 5 μm were prepared using a microtome and used for histological and immunohistochemical staining. Safranin O/Fast Green staining was performed to evaluate cartilage-like matrix deposition and tissue morphology in the repair region. Picrosirius red staining, with Alcian blue counterstaining, was performed to assess collagen fiber deposition and organization under polarized light microscopy.

For immunohistochemical staining, paraffin sections were deparaffinized, rehydrated, and treated with 3% hydrogen peroxide to block endogenous peroxidase activity. The sections were then permeabilized with Triton X-100 and blocked with 1% bovine serum albumin (BSA). Subsequently, the sections were incubated with primary antibodies against collagen type I and collagen type II (1:300; Novus Biologicals, Centennial, CO, USA) for 1 h, followed by incubation with horseradish peroxidase-conjugated secondary antibodies (1:300; Novus Biologicals) for 1 h. Immunoreactivity was visualized using DAB substrate, and the reaction was stopped by rinsing with distilled water. The sections were counterstained with hematoxylin for 10–20 s, dehydrated through graded ethanol solutions, cleared in xylene, and coverslipped with mounting medium. All stained sections were scanned using a high-resolution slide scanner (NanoZoomer, Hamamatsu, Japan).

### 2.11 Statistical analysis

Quantitative data are presented as mean ± standard deviation. In vitro experiments were performed with at least three independent replicates unless otherwise stated, and animal experiments were performed with n = 5 rats per group. Statistical analyses were performed using GraphPad Prism 10. Comparisons between two groups were analyzed using unpaired two-tailed t-tests. Comparisons among multiple groups were analyzed using one-way ANOVA followed by Tukey’s post hoc test when data met assumptions of normality and homogeneity of variance. A value of p < 0.05 was considered statistically significant.

## Result

In this study, we investigated the effects of PEGDA concentration on the mechanical properties, BMSC-associated matrix deposition, and osteochondral repair performance of PEGDA/HAMA hydrogels. The 7.5% PEGDA/HAMA hydrogel exhibited higher stiffness than the 3.75% formulation and was associated with more evident matrix staining in vitro and improved defect filling in vivo.

### 3.1 Fabrication and characterization of PEGDA/HAMA hydrogel scaffolds

PEGDA was incorporated into the HAMA hydrogel precursor solution, and LAP was used as the photoinitiator to enable blue-light-mediated photocrosslinking, resulting in the formation of PEGDA/HAMA hydrogels (Fig. 1A). After photocrosslinking, both the 3.75% and 7.5% PEGDA/HAMA hydrogels maintained their predefined shapes when formed in molds, indicating suitable shape adaptability and structural stability (Fig. 1B). In addition, the stiff hydrogel maintained its structural integrity during qualitative scalpel-contact handling, suggesting improved resistance to local mechanical deformation (Fig. 1C). Mechanical testing showed that the 7.5% PEGDA/HAMA hydrogel exhibited higher compressive stiffness than the 3.75% PEGDA/HAMA hydrogel. The elastic modulus and compressive stress– strain curves indicated that increasing PEGDA content enhanced the mechanical strength of the hydrogel network (Fig. 1D,E). This result is consistent with the increased polymer content and crosslinking density expected in the high-PEGDA formulation. ¹ H NMR analysis was performed to confirm the successful modification of HA. Characteristic proton signals associated with methacrylate groups were observed at approximately 1.75–2.0 ppm, 4.25–4.5 ppm, and 5.5–5.75 ppm, indicating successful methacrylation of HA. The degree of methacrylation was calculated to be approximately 29% based on peak-area analysis (Fig. 1F).

**Figure 1.**
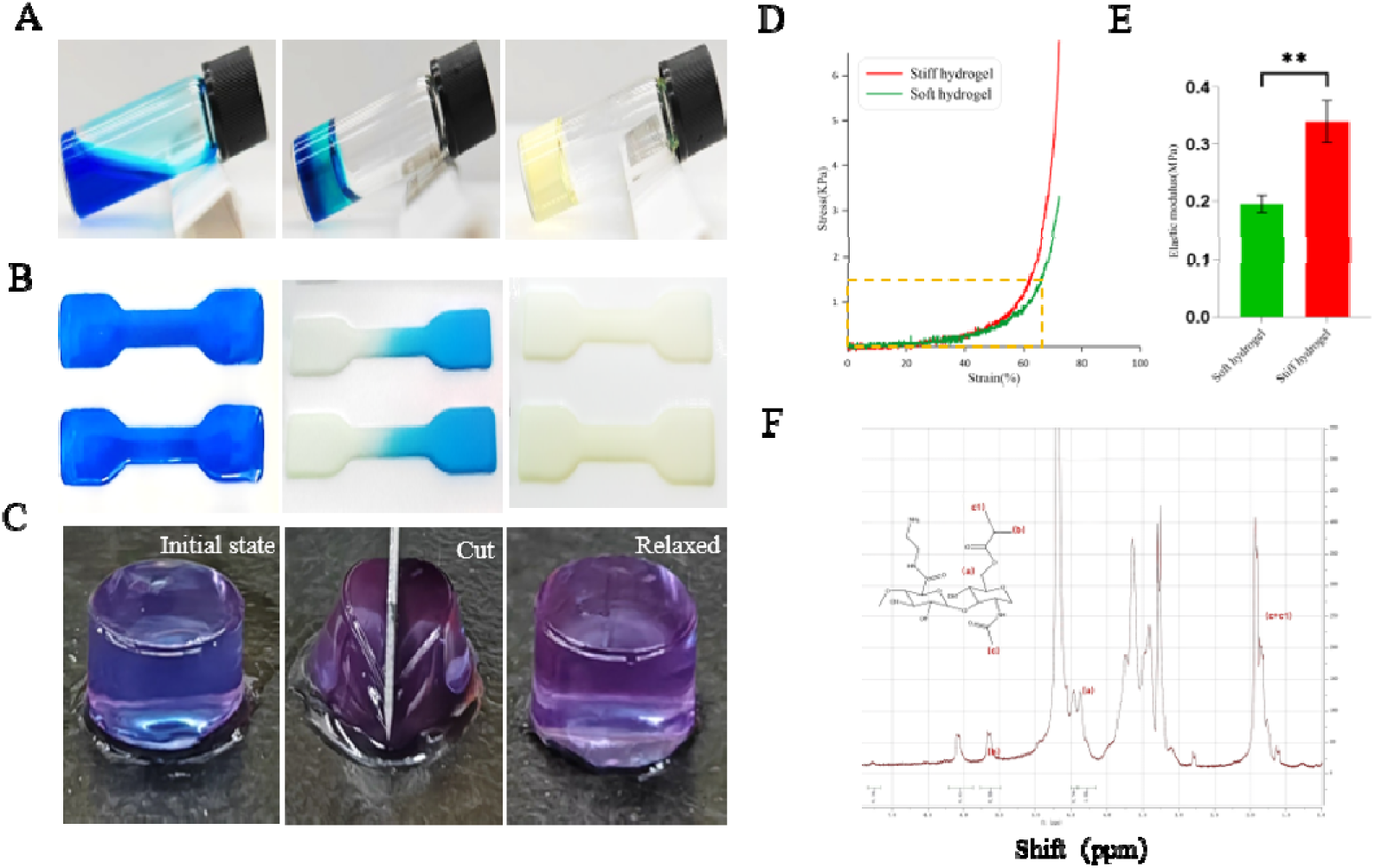
Preparation and characterization of PEGDA/HAMA hydrogels with distinct stiffness profiles. (A) Photographs of PEGDA/HAMA prepolymer solutions after blue-light irradiation. (B) The prepolymer solution was injected into molds and subsequently photocrosslinked to evaluate shape adaptability. (C) Representative photographs showing the qualitative resistance of the photocrosslinked stiff PEGDA/HAMA hydrogel to scalpel-induced deformation, indicating its handling stability and mechanical robustness. (D,E) Elastic modulus and compressive stress–strain curves of the soft hydrogel and stiff hydrogel. (F) ¹H NMR spectrum of methacrylated hyaluronic acid (HAMA).

### 3.2 Surface morphology of BMSC-laden PEGDA/HAMA hydrogels

SEM was used to observe the microstructure of BMSC-laden PEGDA/HAMA hydrogels after lyophilization. Both the soft and stiff hydrogels exhibited interconnected porous structures (Fig. 2A). Compared with the soft hydrogel, the stiff hydrogel showed a relatively more uniform porous architecture with smaller pore sizes. Cell-like structures consistent with encapsulated BMSCs were observed within the hydrogel matrix and are marked in the images. These results suggest that the PEGDA/HAMA hydrogels provided a porous microenvironment suitable for BMSC encapsulation.

**Figure 2.**
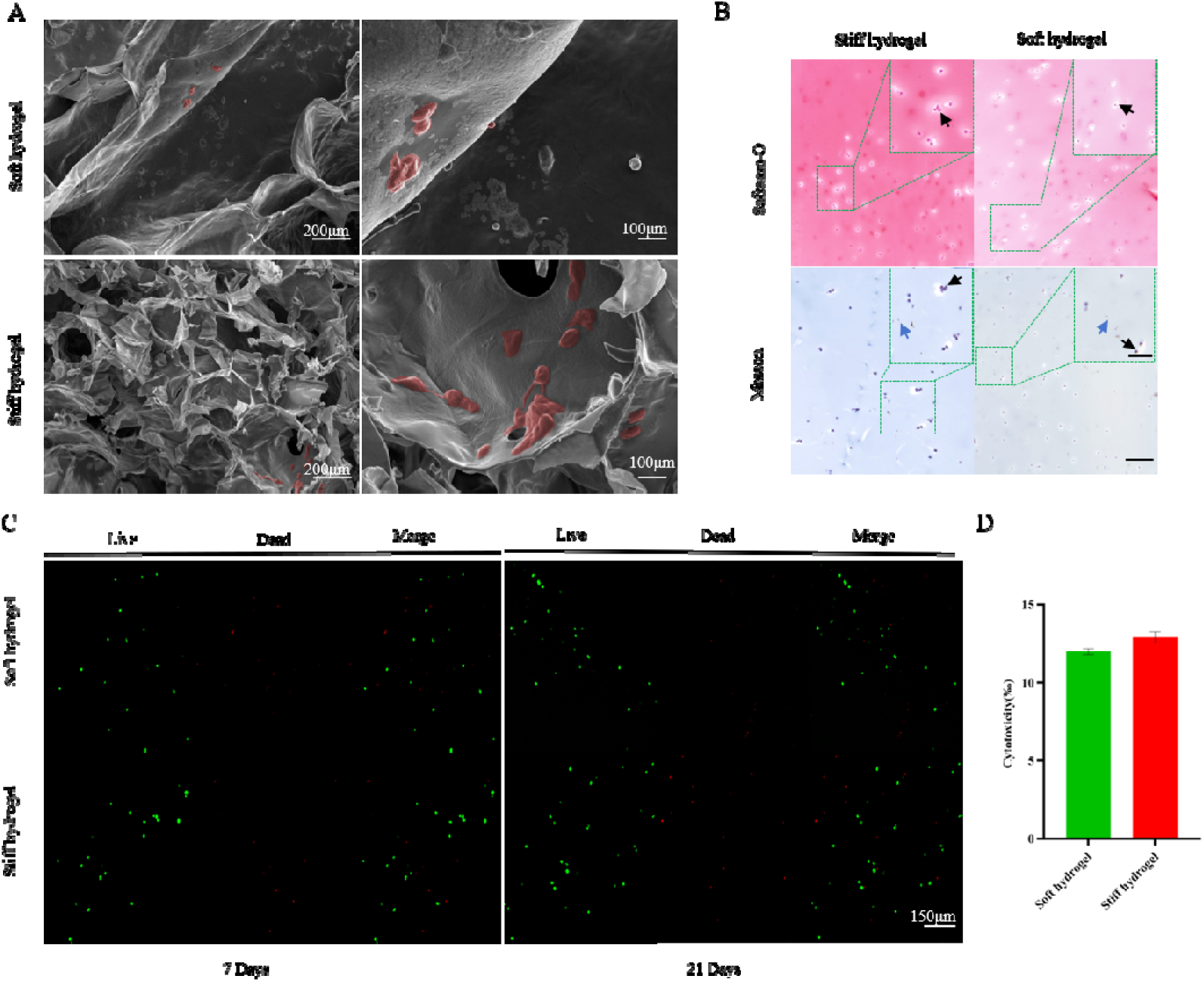
Hydrogel morphology, chondrogenic matrix deposition, and cytocompatibility. (A) SEM images of BMSC-laden soft and stiff PEGDA/HAMA hydrogels. Burgundy pseudo-coloring highlights representative cell-like structures consistent with encapsulated BMSCs. (B) Safranin O and Masson’s staining of BMSC-laden hydrogels shows chondrogenic matrix deposition. Arrows point to cells (black) and residual hydrogel (blue). Scale bars = 200 μm, 100 μm. (C) Live/dead staining of BMSCs encapsulated in PEGDA/HAMA hydrogels at days 7 and 21. Green fluorescence indicates live cells, and red fluorescence indicates dead cells. (D) Quantitative analysis of cytotoxicity based on live/dead staining. Results are representative of three independent experiments.

### 3.3 Chondrogenic matrix deposition of BMSCs in PEGDA/HAMA hydrogels

Chondrogenic induction experiments were performed using BMSCs encapsulated in PEGDA/HAMA hydrogels with different PEGDA concentrations. After a total culture period of 14 days, including 7 days of initial culture and 7 days of chondrogenic induction, matrix deposition was evaluated by Safranin O and Masson’s trichrome staining (Fig. 2B). BMSCs encapsulated in the stiff hydrogel showed more evident Safranin O staining and Masson’s trichrome staining than those in the soft hydrogel, suggesting increased early matrix deposition. However, these staining results should be interpreted as evidence of early chondrogenic matrix accumulation rather than definitive confirmation of mature hyaline cartilage matrix formation.

### 3.4 Cytocompatibility of PEGDA/HAMA Hydrogels

The cytocompatibility of PEGDA/HAMA hydrogels was assessed using live/dead staining. Live/dead images showed that most encapsulated BMSCs remained viable in both hydrogel formulations after 7 and 21 days of culture. Quantitative analysis showed low cytotoxicity in both the soft and stiff hydrogel groups, with no significant difference between the two formulations at either time point (Fig. 2C,D). These results indicate that both PEGDA/HAMA hydrogels exhibited acceptable cytocompatibility toward BMSCs.

### 3.5 Gross evaluation of osteochondral defect repair in rats

To evaluate the efficacy of PEGDA/HAMA hydrogels in osteochondral defect repair in vivo, a rat model with articular cartilage defects was employed. The subjects were randomly assigned to three groups: a defect-only group, a soft hydrogel group, and a stiff hydrogel group. At 4 weeks post-operation, gross inspection and ICRS scoring [28] revealed that all groups except the defect-only group showed partial defect filling. By 8 weeks, the stiff hydrogel group showed signs of repair with a less distinct boundary between the regenerated tissue and adjacent normal tissue. The defect-only group showed limited regeneration, whereas the soft hydrogel group demonstrated partial repair tissue formation (Fig. 3).

**Figure 3.**
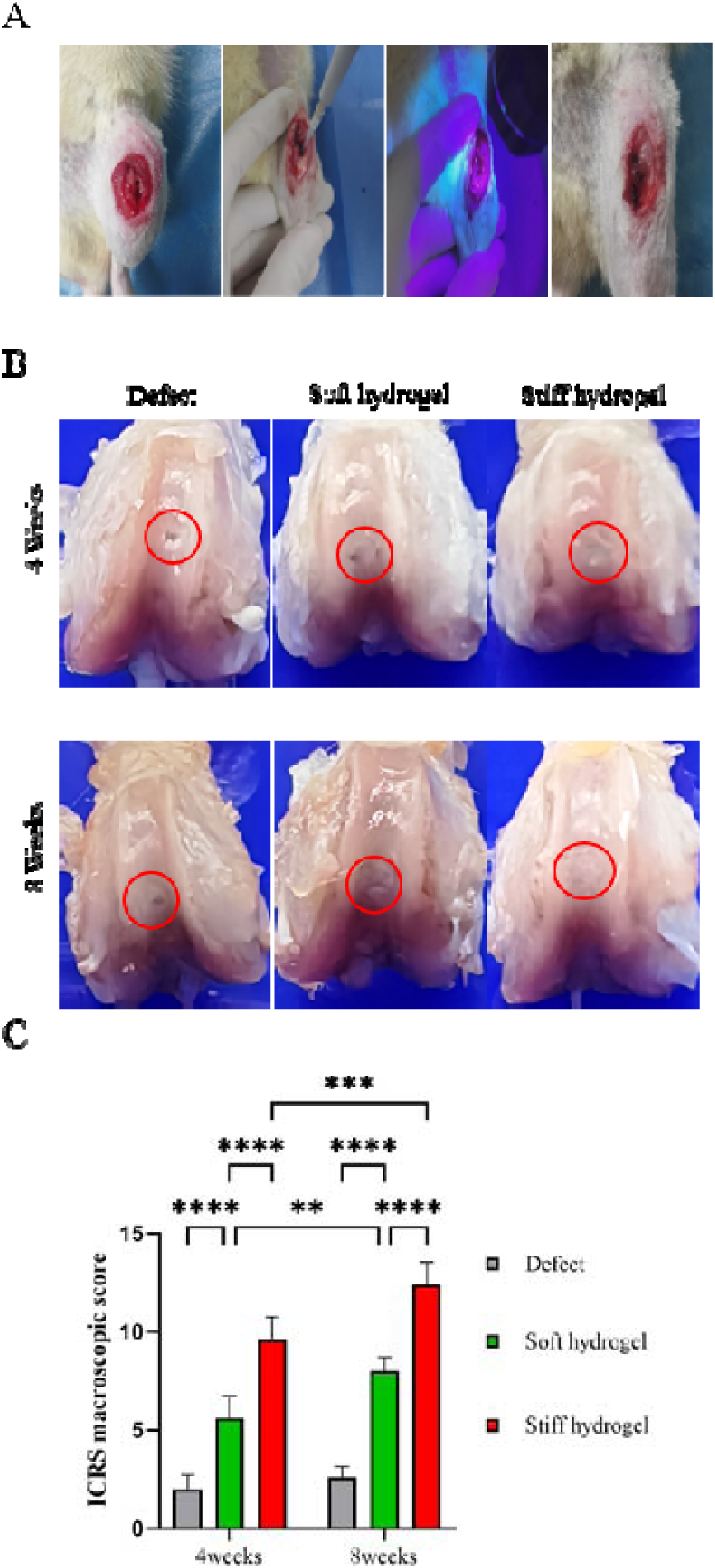
Gross evaluation of osteochondral defect repair at 4 and 8 weeks after implantation. Stiff hydrogel: BMSC-laden 7.5% PEGDA/HAMA hydrogel; Soft hydrogel: BMSC-laden 3.75% PEGDA/HAMA hydrogel. (A) Schematic illustration of the osteochondral defect procedure. (B) Gross images of the Defect, Soft hydrogel, and Stiff hydrogel groups at 4 and 8 weeks post-surgery. Red circles indicate the original defect region. (C) Quantitative analysis based on ICRS criteria at 4 and 8 weeks post-surgery. Data are presented as mean ± s.d. (n = 5).

### 3.6 In vivo evaluation of osteochondral defect repair

As previously described, the femurs from the treated rats were collected at 4 and 8 weeks post-surgery for micro-CT evaluation. Results showed that the subchondral bone surfaces in the Stiff hydrogel group were smooth, uniform, and continuous, with no significant collapse observed at 8 weeks. In contrast, the other groups exhibited rougher subchondral surfaces with varying degrees of damage. The stiff hydrogel group showed higher BV/TV, Tb.N, and Tb.Th, suggesting improved subchondral bone remodeling within the defect region. Meanwhile, the soft hydrogel group showed higher Tb.Sp, suggesting less favorable subchondral bone remodeling compared with the stiff hydrogel group. (Fig. 4).

**Figure 4.**
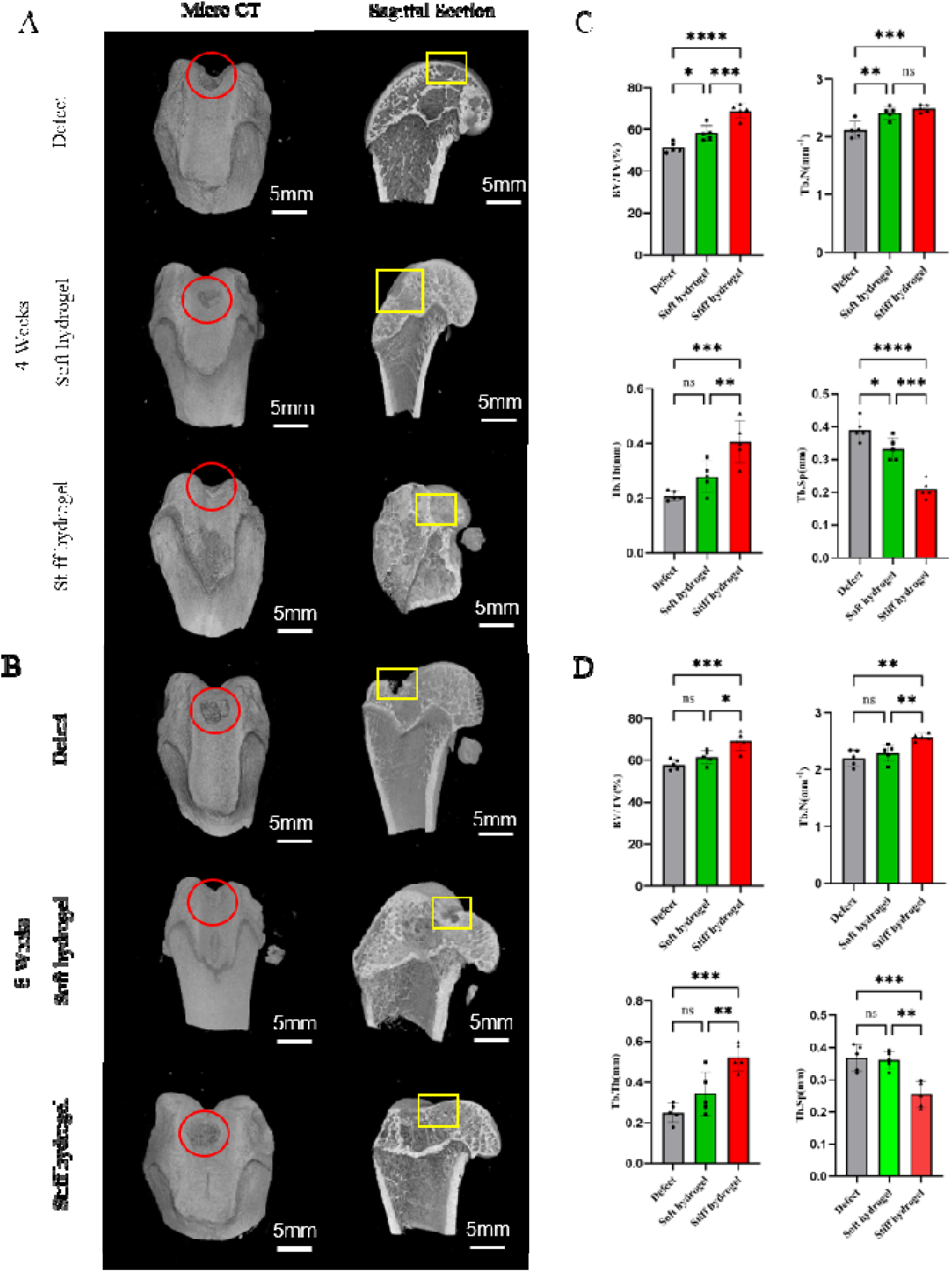
Micro-CT evaluation of subchondral bone remodeling in the rat osteochondral defect model. (A,B) Micro-CT images of the Defect, Soft hydrogel, and Stiff hydrogel groups at 4 and 8 weeks post-surgery. Red circles indicate the defect region in the anterior view, and yellow boxes indicate the defect region in the sagittal view. (C,D) Quantitative analysis of subchondral bone parameters, including BV/TV, Tb.N, Tb.Sp, and Tb.Th. Data are presented as mean ± s.d. (n = 5). Statistical significance is indicated as *p < 0.05, **p < 0.01, and ***p < 0.001; ns indicates no significant difference.

### 3.7 Histological and immunohistochemical staining

Safranin O/Fast Green staining was performed to evaluate matrix deposition and tissue morphology in the repair region (Fig. 5). At 4 weeks, the defect group showed limited tissue filling, whereas the hydrogel-treated groups exhibited partial defect filling. The stiff hydrogel group showed relatively improved tissue continuity compared with the soft hydrogel group. At 8 weeks, the stiff hydrogel group demonstrated more complete defect filling and better integration with adjacent tissue than the defect and soft hydrogel groups. However, the newly formed tissue did not fully resemble native hyaline cartilage.

**Figure 5.**
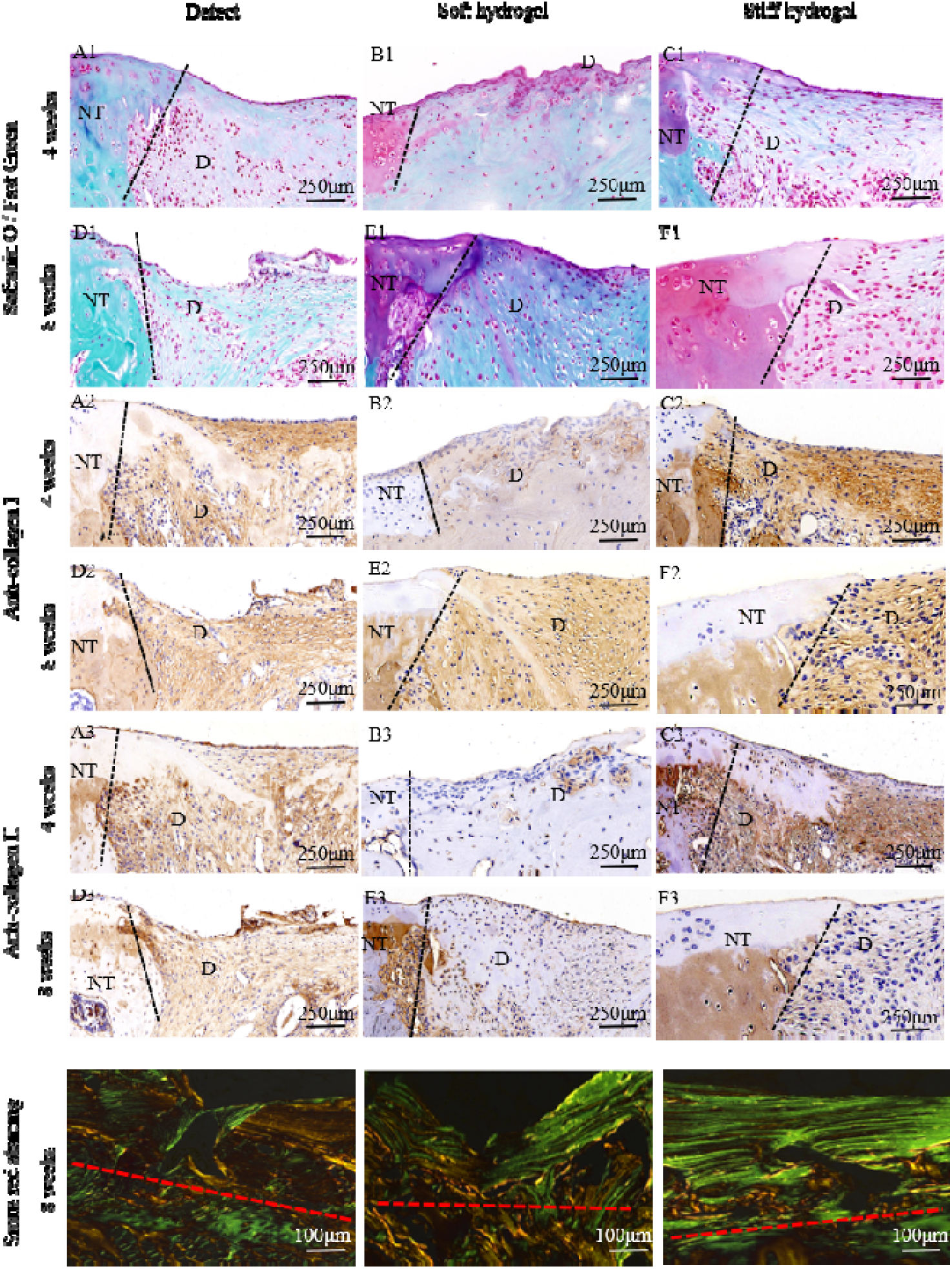
Histological and immunohistochemical evaluation of osteochondral repair at 4 and 8 weeks post-surgery. A–C represent 4-week samples from the Defect, Soft hydrogel, and Stiff hydrogel groups, respectively; D–F represent 8-week samples from the Defect, Soft hydrogel, and Stiff hydrogel groups, respectively. From top to bottom, the first three rows show Safranin O/Fast Green staining, collagen type I immunohistochemistry, and collagen type II immunohistochemistry, respectively. Sirius red staining under polarized light was used to evaluate collagen fiber deposition and organization at 8 weeks. D indicates the defect region, NT indicates native tissue, and the dotted line indicates the boundary of newly formed tissue.

Immunohistochemical staining further showed that collagen type I was predominantly deposited in the repair region, particularly in the hydrogel-treated groups. In contrast, collagen type II staining was mainly observed in the surrounding native cartilage, with limited staining in the central repair tissue. These results indicate that the regenerated tissue was primarily fibrocartilaginous rather than hyaline-like cartilage. Sirius red staining under polarized light further showed collagen fiber deposition in the repair area, supporting the presence of collagen-rich repair tissue but not confirming restoration of native hyaline cartilage architecture.

## Discussion

In this study, we developed a BMSC-laden PEGDA/HAMA dual-crosslinked hydrogel system with distinct stiffness profiles for osteochondral defect repair. By varying the PEGDA concentration from 3.75% to 7.5%, we established a binary stiffness-based comparison to evaluate how PEGDA-content-dependent mechanical reinforcement influences BMSC-associated matrix deposition and early osteochondral repair outcomes. The main finding of this study is that increased PEGDA content enhanced compressive stiffness, supported early matrix deposition in vitro, and improved defect filling and subchondral bone remodeling in vivo. Importantly, the present results also show that stiffness modulation alone was insufficient to restore a collagen type II-rich hyaline-like cartilage matrix within the 8-week observation period.

The combination of covalent crosslinking and non-covalent interactions may have contributed to the mechanical stability and structural integrity of the PEGDA/HAMA hydrogel network, consistent with previous studies showing that hydrogel network composition can influence matrix properties and encapsulated MSC behavior [29]. In the present study, compressive mechanical testing indicated that the 7.5% PEGDA/HAMA hydrogel exhibited higher mechanical strength than the 3.75% formulation. This enhanced stiffness may provide greater early structural support within the defect region, which is particularly important in a mechanically active joint environment. The HAMA component was prepared by introducing methacrylate groups onto HA using MMA, and successful modification was confirmed by ¹H NMR. Live/dead staining further showed that both hydrogel formulations exhibited acceptable cytocompatibility toward BMSCs, suggesting that the increased PEGDA content used in the stiff formulation did not induce obvious cytotoxicity under the present experimental conditions.

The microstructure of hydrogels is closely associated with their biological performance. SEM imaging revealed that the stiff hydrogel containing 7.5% PEGDA possessed a relatively uniform porous structure with smaller pore sizes of approximately 50 μm. Previous studies have shown that scaffold pore size can influence MSC proliferation, chondrogenic gene expression, and cartilage matrix synthesis [30]. In this study, the porous architecture of the stiff hydrogel may partially explain the more evident Safranin O and Masson’s trichrome staining observed in vitro and the improved defect filling observed in vivo. However, because only two PEGDA concentrations were compared, these findings should be interpreted as a binary stiffness comparison rather than a complete demonstration of continuous mechanical tunability.

Both hydrogels supported BMSC survival in vitro, but the biological responses appeared to differ between the two stiffness conditions. The stiff hydrogel supported more evident early chondrogenic matrix deposition by BMSCs, as indicated by Safranin O and Masson’s trichrome staining. Nevertheless, these staining results should be interpreted as evidence of early matrix accumulation rather than definitive proof of mature hyaline cartilage formation. This interpretation is important because matrix staining alone cannot fully demonstrate stable chondrogenic phenotype acquisition or hyaline cartilage-specific matrix formation.

In vivo, BMSC-laden PEGDA/HAMA hydrogels with different PEGDA concentrations were implanted into rat osteochondral defects. At 8 weeks, the 7.5% PEGDA/HAMA hydrogel showed improved defect filling and subchondral bone remodeling compared with the 3.75% PEGDA/HAMA hydrogel and defect groups, as shown by gross observation, histological staining, and micro-CT analysis. These findings suggest that increasing PEGDA content may enhance the early structural support provided by the hydrogel, thereby contributing to improved early osteochondral defect repair.

Nevertheless, the histological and immunohistochemical findings indicate that the repair tissue was mainly fibrocartilaginous. Collagen type I was predominantly detected in the repair region, whereas collagen type II expression remained limited and was mainly localized to the surrounding native cartilage. Therefore, the present data do not support complete hyaline cartilage regeneration. Instead, they suggest that the stiff BMSC-laden PEGDA/HAMA hydrogel improved early defect filling and subchondral bone remodeling but was insufficient to restore a hyaline-like cartilage matrix within the 8-week observation period.

Recent studies have increasingly emphasized that hydrogel-based osteochondral repair requires not only mechanical support but also appropriate biological and architectural cues. For example, Pei et al. developed an exosome-functionalized GelMA/CSMA/HAMA hydrogel incorporating BMSC-derived exosomes and reported enhanced BMSC recruitment, chondrogenic differentiation, and hyaline-like cartilage regeneration in a rat osteochondral defect model [31]. In contrast, the present PEGDA/HAMA system did not include exosomes, growth factors, or other chondroinductive biological cues, which may partially explain the limited collagen type II deposition observed in the repair region. Similarly, recent dual-layer or zonal hydrogel scaffolds have attempted to mimic the native osteochondral interface by spatially separating cartilage- and bone-supportive microenvironments. Zhan et al. reported a DLP-printed dual-layer SilMA-based scaffold incorporating PEGDA, HAp, and chondrocyte-laden microspheres for osteochondral defect reconstruction [32], while Cao et al. developed an injectable bilayer collagen/chondroitin sulfate hydrogel scaffold for osteochondral repair [33]. Compared with these biomimetic multilayered systems, our PEGDA/HAMA hydrogel used a simpler single-phase formulation and therefore primarily provided early structural support rather than a spatially organized cartilage–bone regenerative microenvironment. These comparisons suggest that increased PEGDA content can improve early defect filling and subchondral bone remodeling, but stiffness modulation alone may be insufficient to drive stable hyaline-like cartilage matrix formation. This interpretation is also supported by recent findings showing that interpenetrating hydrogel network composition can regulate encapsulated MSC spreading and in vivo degradation behavior [34].

Despite the absence of complete hyaline cartilage regeneration, the present study provides useful information for the design of hydrogel-based osteochondral repair systems. First, it demonstrates that increasing PEGDA content in a BMSC-laden PEGDA/HAMA hydrogel can enhance compressive stiffness and improve early osteochondral defect filling. Second, it shows that the stiffer hydrogel formulation is more favorable for subchondral bone remodeling than the softer formulation in a rat defect model. Third, the limited collagen type II expression observed in vivo highlights an important design limitation: mechanical reinforcement alone does not necessarily translate into hyaline cartilage matrix restoration. Therefore, the main contribution of this study is not the achievement of complete cartilage regeneration, but the identification of a stiffness-dependent structural support effect in BMSC-laden PEGDA/HAMA hydrogels and the clarification that additional biological or zonal cues are required for hyaline cartilage-oriented repair.

Several factors may have contributed to the fibrocartilaginous repair outcome. First, the 8-week observation period may not have been sufficient for complete matrix maturation and remodeling. Previous osteochondral scaffold studies have suggested that scaffold retention and prolonged degradation may influence histological interpretation and that longer follow-up periods are required to fully evaluate repair outcomes [35]. Second, the degradation behavior of the PEGDA/HAMA hydrogel was not directly quantified in this study; therefore, any relationship between scaffold persistence and matrix remodeling should be interpreted cautiously. Previous PEGDA/HA network studies have also shown that hydrogel composition and cell-mediated degradation can influence three-dimensional cell spreading and migration [36]. Third, the complex post-operative environment in the rat joint may have affected matrix remodeling and repair tissue phenotype. In addition, the importance of cartilage zonal organization has been emphasized in recent stratified cartilage repair strategies [37]. Further studies incorporating longer observation periods, quantitative degradation analysis, and more detailed matrix characterization are required to optimize this hydrogel system for hyaline cartilage-oriented osteochondral repair.

This study has several limitations. First, only two PEGDA concentrations were evaluated; therefore, the present findings should be interpreted as a binary stiffness comparison rather than a complete concentration-gradient analysis. Additional intermediate PEGDA concentrations are required to establish a more comprehensive concentration–structure–function relationship. Second, the observation period was limited to 8 weeks, which may be insufficient to evaluate long-term matrix maturation, hydrogel remodeling, and the final quality of the repaired tissue. Third, hydrogel degradation kinetics were not directly quantified, and the relationship between scaffold persistence, tissue ingrowth, and collagen matrix remodeling remains unclear. Finally, although the 7.5% PEGDA/HAMA hydrogel improved defect filling and subchondral bone remodeling, the predominance of collagen type I and limited collagen type II expression indicate that the repair tissue remained mainly fibrocartilaginous. Future studies should incorporate longer observation periods, quantitative degradation analysis, additional stiffness gradients, and more detailed matrix characterization to optimize this BMSC-laden PEGDA/HAMA hydrogel system for hyaline cartilage-oriented osteochondral repair.

## Conclusion

In conclusion, this study developed BMSC-laden PEGDA/HAMA hydrogels with two representative stiffness levels and evaluated their effects on osteochondral defect repair. The 7.5% PEGDA/HAMA hydrogel exhibited higher stiffness than the 3.75% formulation and supported BMSC viability and matrix deposition in vitro. In the rat osteochondral defect model, the stiff hydrogel improved defect filling and subchondral bone remodeling compared with the soft hydrogel and defect groups. However, the repair tissue showed predominant collagen type I deposition and limited collagen type II expression, indicating fibrocartilaginous rather than hyaline-like cartilage repair. These findings suggest that PEGDA/HAMA hydrogels may serve as a useful platform for osteochondral repair, but further optimization of degradation behavior, matrix maturation, and collagen type II-rich matrix deposition is required.

## Author Contributions

Conceptualization, M.J.L., X.T.Z. and Y.H.H.; methodology, X.G. and L.Q.N.; investigation, Y.S. and J.J.L.; data curation, J.J.L., M.J.L. and Y.S.; writing—original draft preparation, L.Q.N., M.J.L. and X.G.; writing—review and editing, M.J.L., X.F.Z. and Y.H.H.; supervision, X.T.Z., X.F.Z. and Y.H.H.; funding acquisition, X.T.Z. and Y.H.H. All authors have read and agreed to the published version of the manuscript.

## Data Availability Statement

The data presented in this study are available from the corresponding author upon reasonable request.

## Institutional Review Board Statement

All animal procedures were approved by the Institutional Animal Care and Use Committee of Peking University Shenzhen Hospital under protocol No. 2023-139, approved on 21 June 2023, and were performed in accordance with institutional guidelines for animal care and use.

## Informed Consent Statement

Not applicable.

